# JN403, an alpha-7-nicotine-acetylcholine-receptor agonist, reduces alpha-synuclein induced inflammatory parameters of *in vitro* microglia but fails to attenuate the reduction of TH positive nigral neurons in a focal alpha-synuclein overexpression mouse model of Parkinson’s disease

**DOI:** 10.1101/2020.04.04.996892

**Authors:** Bolam Lee, Carmen Noelker, Dominik Feuerbach, Lars Timmermann, W.-H Chiu, Wolfgang H. Oertel

## Abstract

Alpha-7-nicotine-acetylcholine-receptor (α7-nAChRs) agonists modulate the cholinergic antiinflammatory pathway to attenuate proinflammatory signals and reduce dopaminergic neuronal cell loss in toxin-induced experimental murine models of Parkinson’s disease (PD). The protein α-synuclein (αSyn) is considered to represent the major pathogenic component in the etiology and progression of sporadic PD. However, no research has been performed to evaluate the effect of α7-nAChR agonists in human αSyn mediated models of PD. We, therefore, investigated the effect of the compound JN403, an α7-nAChR specific agonist, in αSyn treated *in vitro* microglia culture and in a human αSyn overexpression *in vivo* mouse model. In primary mouse microglia cells, αSyn fragment 61-140 treatment increased the release of nitric oxide (NO), tumor necrosis factor (TNF)-α and interleukin (IL)-6, and decreased cell viability. In contrast, 100 nM or 1 μM of JN403 co-incubation significantly reduced the level of NO and TNF-α release in the microglial cells. For in-vivo testing of JN403, a recombinant adeno-associated viral vector (rAAV)-mediated unilateral intranigral overexpression of human wild-type-αSyn (WT-αSyn) or of the control protein luciferase (luc) was induced via stereotactic delivery in C57/BL6N mice. Targeted WT-αSyn overexpression reduced 20% of the number of tyrosine hydroxylase (TH) positive (+) nigral neurons after 10 weeks. Subcutaneous daily treatment of 30 mg/kg JN403 over 9 weeks starting at postoperative week 1 did not alter the decrease of TH+ neuronal numbers, and microglial density in WT-αSyn overexpression mouse model. The reduced density of TH+ striatal terminals in the WT-αSyn groups was also not recovered by the JN403 treatment. In summary, JN403, an α7-nAChR specific agonist shows a beneficial effect on ameliorating proinflammatory signals in αSyn exposed microglia cells. However, no significant in-vivo treatment effect was found in an intranigral WT-αSyn overexpression mouse model of PD.

## INTRODUCTION

Epidemiological studies have repeatedly shown a robust inverse relationship between smoking and a reduced risk of developing Parkinson’s disease (PD) [1,2]. Nicotine is a major component of tobacco and easily reaches the CNS during smoking. Thus the epidemiologic data suggest that exposure of the body including the nervous system to nicotine for many years during the prodromal stages of PD (Braak stage 1-3) [3], i.e. before the onset of the cardinal motor signs of akinesia, tremor at rest or rigidity, may be beneficial and may delay the time to or even prevent conversion to “manifest” PD [4–6]. In contrast, it is conceivable that nicotine has no beneficial effect on manifest PD and accordingly clinical studies probing therapeutic efficacy of nicotine in de novo and advanced PD patients (Braak stage 4 and higher) did not show a convincing symptomatic (improvement of motor symptoms) or disease-modifying effect [8–11].

This inconsistent result may imply a certain specific role of nicotine in each neuropathologic stage of PD. One possible mechanism of how nicotine may act is the ‘cholinergic anti-inflammatory pathway’. Substantial experimental studies with nicotinic agonists demonstrate that α-7-nicotine-acetylcholine-receptors (α7-nAChRs) of the AChR-family are pivotal to inhibit the synthesis and release of proinflammatory cytokines, hence treatment with nicotinic agonists may modulate the production of proinflammatory cytokines from immune cells [12–14]. Chronically activated microglia release abundant proinflammatory mediators, such as nitric oxide (NO), tumor necrosis factor-α (TNF-α), and reactive oxygen species, which in turn may trigger further microglial activation and neuronal loss [15,16].

Neuroinflammation is one of the major characteristics of PD pathology [17], however, a precise role of neuroinflammation, especially in regards to gliosis and neurodegeneration, still needs to be uncovered. There are now several lines of evidence suggesting that murine primary cultured microglial cells express the α7-nAChR subunit, and the microglial release of TNF-α and IL-6 was modulated with acetylcholine or nicotine treatment when challenged by lipopolysaccharide (LPS) via activation of α7-nAChRs [18,19]. Besides, α7-nAChRs play a critical role in “neuroprotection” against MPTP intoxication as shown in the α7-nAChR KO transgenic mouse line [20] as well as in our previous work proposing that MPTP-induced loss of nigral neurons in mice was prevented by treatment with the selective α7 nACh agonist, PNU-282987 [21]. Therefore, α7-nAChRs could serve as a crucial link between inflammation and neurodegeneration [21,22] and may represent a pharmacological target for potential neuroprotective therapeutic strategies.

The protein α-synuclein (αSyn) is considered to represent the major pathogenic component in the etiology and progression of sporadic PD and is composed of 140 amino acid containing three functional domains: the N-terminal domain amino acid residues (AA 1-60), the NAC-region (AA 1-95) and the C-terminal domain (AA 96-140) [23,24]. αSyn can self-assemble into oligomers and fibrils, and it has been extensively used with different genetic modifications and aggregation forms over the years to model progressive pathological conditions in experimental settings [25]. However, α7-nAChR agonists in human αSyn mediated models have rarely been investigated yet. To the extent of our knowledge, there is a single report investigating the effect of nicotine on a transgenic mouse model overexpressing wild type human αSyn under the Thy1-promoter (Thy 1 -αSyn mice). In this report, long-term subcutaneous treatment of nicotine did not diminish αSyn aggregation, nor microglial activation [26]. Since this Thy1-αSyn mouse strain did not show a significant loss of dopaminergic neurons in the substantia nigra pars compacta (SNc), it is difficult to interpret a potential neuroprotective effect of nicotine in this model. On the contrary, the viral-mediated overexpression of αSyn has been established and extensively characterized for more than a decade [27]. It has been studied as a common model to study the αSyn toxicity in the SNc including ours [28], in this study, we proposed a neuroprotective effect of a phosphodiesterase 1 inhibitor using a viral vector-mediated wild-type αSyn overexpression. Moreover, other PD-vulnerable neuronal populations such as the locus coeruleus [29] and the dorsal motor nucleus of the vagal nerve [30] have been investigated using the viral-mediated overexpression model.

Based on our previous experiences, we investigated JN403, a novel α7-nAChRs agonist using the same in-vivo experimental scheme. Previous research findings with JN403 suggest that this compound shows good penetration into the brain after oral administration in rodents and may be beneficial to alleviate mechanical sensory pain and anxiety [31,32]. These observations led to the questions, whether JN403 has the potential to modulate *in vitro* microglial function and display efficacy in a viral vector-mediated αSyn overexpression rodent model for PD. We, therefore, studied the effect of the JN403 treatment on neuroinflammatory signals in microglial cell cultures incubated with a human αSyn fragment as well as on the loss of dopaminergic neurons in an αSyn overexpression *in vivo* mouse model after unilateral intranigral injection of an AAV vector carrying the gene for human wild type αSyn.

## 1. MATERIAL AND METHODS

### 1.1. Microglial cultures of the mouse ventral mesencephalon

The protocol of the microglial culture has been published [24,33]. Briefly, microglial cultures derived from E13.5 mouse embryos (Janvier Breeding Center, France) were obtained by mild trypsinization as described previously. After 14 days, cultures were washed with Dulbecco’s modified Eagle’s medium/F-12 nutrient mixture (DMEM/F12; Invitrogen, Cergy-Pontoise, France) supplemented with 10% fetal calf serum (FCS) and then incubated with diluted DMEM/F12 (trypsin) 1:4 for 90 min at 37 °C until the astrocytic upper layer was detached. The medium containing the layer of detached cells was aspirated, and the highly enriched microglial population that remained attached to the bottom of the well was exposed to 500 μL of DMEM/F12 with 10% FCS to allow trypsin inactivation. The remaining microglial cells were treated with JN403 (100 nM, 1 μM or 10 μM) or control for 24 hours followed by addition of human α-synuclein (αSyn) fragment (61-140) at 1 μM (rPeptide; Bogart, GA, USA) for another 24 hours. This αSyn fragment and its concentration were carefully selected as it induced the most marked toxicity in our previous experimental setting [24]. With this (αSyn) fragment we also observed an increased aggregation tendency at the C-terminal domain including the NAC region of AA 61-95. Furthermore, this observation is in alignment with in-vitro results reported by other groups [34,35]. All *in vitro* experiments were performed using the same microglial cultured cells described above with a minimum of 3 wells per experimental condition. The same procedures were repeated at 4-5 different trials and the data were pooled afterward to analyze.

### 1.2. MTT-assay

The viability of microglial cells was measured using the 3-(4,5-dimethylthiazol-2-yl)-2,5-diphenyltetrazolium bromide (MTT; Sigma-Aldrich, Steinheim, Germany) assay as described previously [36]. MTT was added to the cells at a final concentration of 0.5 mg/ml in the cell culture medium, and the cells were incubated for 30 min at 37 °C. Afterward, cells were lysed using Dimethyl sulfoxide (DMSO) (AppliChem; Gatersleben, Germany) and the absorptions from each well were taken at 570 nm in the dark using a microplate reader (Infinite M200, Tecan, Crailsheim, Germany).

### 1.3. Measurement of nitric oxide (NO)

Accumulated nitrite, a stable oxidation product of NO, was measured using Griess reagents. 50 μL of primary microglial cell supernatant was transferred to a 96-well microtiter plate, and 50 μL of 1% sulfanilamide in 5% phosphoric acid was added. After 10 minutes of incubation in the dark, 50 μL of 0.1% naphthyl ethylenediamine dihydrochloride was added followed by further incubation for an additional 10 minutes in the dark. The absorbance was measured at 450 nm using a plate reader (ELISA-Reader Infinite^®^ 200 series, Tecan; Crailsheim, Germany).

### 1.4. Cytokine quantification by ELISA

Microglial cells were collected and the release of the proinflammatory cytokines (TNF-α and IL-6) in the supernatant was measured by using the DuoSet ELISA Development System with mouse TNF-α and IL-6 (e-bioscience; Darmstadt, Germany). The ELISAs were carried out according to the manufacturer’s protocol. The initial level of cytokine release (TNF-α and IL-6) was assessed with a single pre-incubation of JN403 (100 nM or 1 μM), and the changes of each cytokine were measured again after 24-hour exposure to human α-synuclein fragment (61-140) - compared to the control-on the following day.

### 1.5. Animals

A total number of 30 male C57/BL6N mice (Charles River; Sulzfeld, Germany), 10 weeks old at the beginning of the experiment, were used. The mice were housed in standard cages with *ad libitum* access to food and water at 23°C with a 12/12 hours light/dark cycle. Animal experiments were performed according to German legislation and mice were handled according to the EU Council Directive 86/609/EEC. Approval of the experiment from the responsible institutional governmental agency (Regierungspräsidium Giessen, Germany) was documented under the reference code V54-19 c 20 15h 01 MR 20/15 Nr. 116/2014.

### 1.6. Targeted αSyn overexpression via the delivery of recombinant AAV5 in the SNc

Mice were deeply anesthetized with a mixture of Ketavet^®^ (100 mg/kg, Zoetis Deutschland GmbH; Berlin, Germany) and 2% Rompun^®^ (5 mg/kg, Bayer; Leverkusen, Germany), diluted in 0.9% NaCl. Later mice were placed in a stereotactic frame (Kopf Instruments, Tujunga, CA) for operation [37]. A total amount of 2μl rAAV5-CBA-human-α-synuclein (wild-type; original titer: 1×10^13^ vg/ml) or rAAV5-CBA-luciferase (original titer: 1×10^13^ vg/ml) was unilaterally injected into right side of the SNc over ten minutes (flow rate of 200 nl/min) by microinjection pump (Micro4TM, WPI, Sarasota, USA). After the injection, the needle stayed in the brain for an additional 10 minutes before it was slowly retracted. The injections were performed by using a microsyringe, stainless steel needle (33G, WPI, Sarasota, USA). The coordinates of the injection were anteroposterior: −3.1 mm, mediolateral: - 1.2 mm, and dorsoventral: −4.2 mm, relative to bregma using a flat skull position [38].

The viral-mediated vectors were provided by the Michael J. Fox Foundation, and the Gene Therapy Center of the University of North Carolina at Chapel Hill, USA.

### 1.7. The *in vivo* experimental design with JN403 treatment

Mice were habituated for 1 week following arrival. To investigate the effects of JN403 on the wild-type human α-synuclein overexpression in the SNc, animals received the injection of either rAAV5-CBA-human-wildtype-α-synuclein (rAAV-WT-αSyn) or rAAV5-CBA-luciferase (rAAV-luc) (as overexpression of a control protein) as previously published [28. Six to eight mice were thereafter assigned to either JN403 treated- or vehicle-treated groups. A total of 4 groups were formed: rAAV-WT-αSyn treated with saline (AN), rAAV-WT-αSyn treated with JN403 (AJ); rAAV-luc treated with saline (LN) and rAAV-luc treated with JN403 (LJ). The dosage of JN403 for subcutaneous injection (s.c) was 30 mg/kg dissolved in saline solution based on the experimental conclusion as previously published [32. One week after the viral injection, mice received a daily s.c injection of JN403 or saline for an additional 9 weeks. Afterward, all mice were sacrificed for further quantification analysis and image processing (**Fig. 1A-B**).

**Figure 1.**
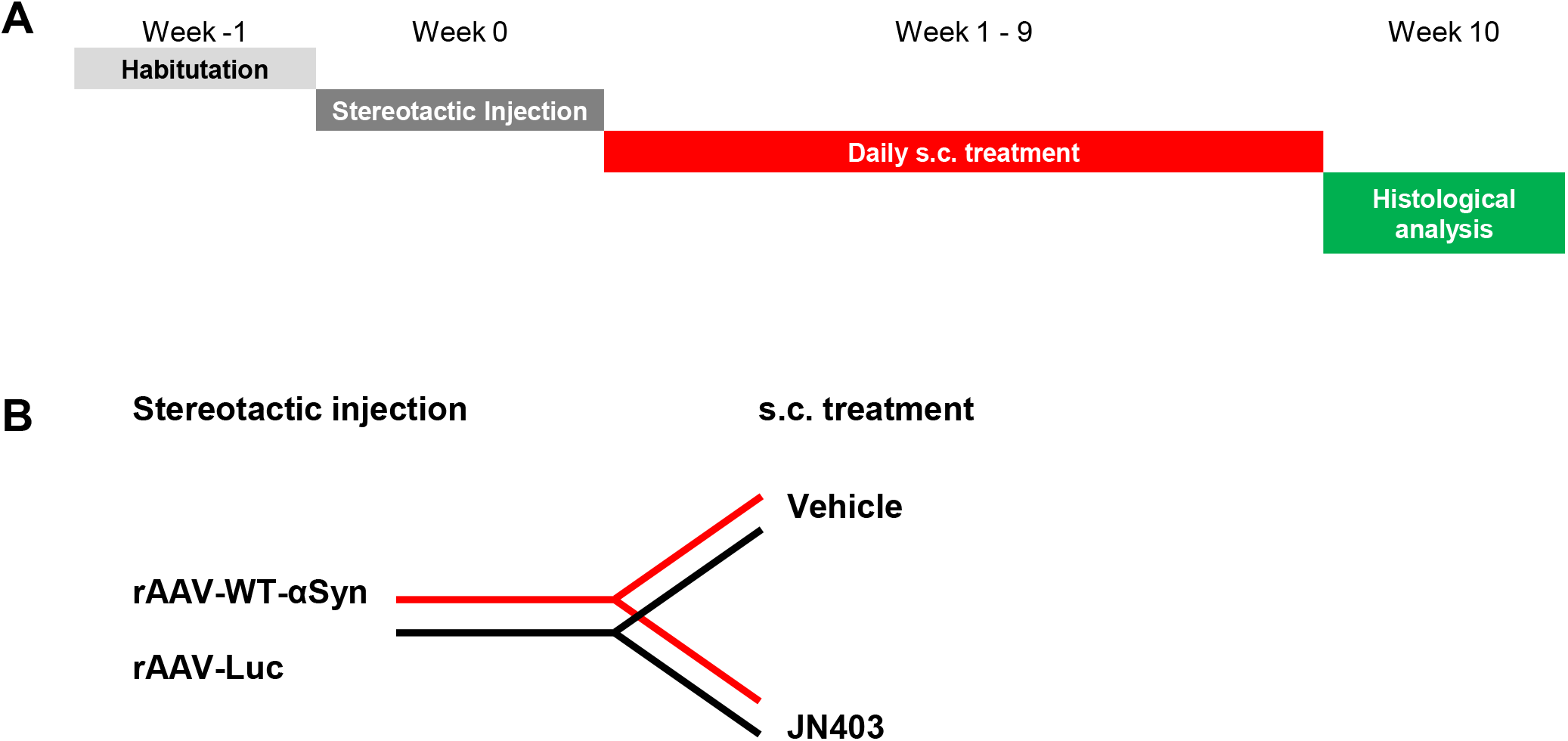
The in-vivo experimental design with JN403 treatment. (A) Mice were habituated for 1 week following arrival, and stereotactic microinjection with viral vectors was performed. To investigate the effects of JN403 mice received a daily s.c injection of JN403 or saline. These injections started 1 week after the operation and were continued for 9 weeks. Afterward, all mice were sacrificed for further quantification analysis and image processing. (B) Mice were operated for the microinjection of the rAAV5-CBA-human-wildtype-α-synuclein (rAAV-WT-αSyn) or rAAV5-CBA-luciferase (rAAV-luc) and administered either with JN403 or vehicle. As a result, 4 different groups were created in this in-vivo experiment.

### 1.8. Tissue preparation and immunohistochemistry

Tissue preparation and the staining protocols were performed as described previously [37]. The animals were deeply anesthetized and perfused transcardially with 0.1 M phosphate-buffered saline (PBS) followed by ice-cold 4% PFA. Coronal sections of the striatum and the midbrain containing the SNc were sliced at 30 μm thickness by a cryostat microtome, collected in 10 regularly spaced series in antifreeze buffer (1:1:3 volume ratio of ethyl glycerol, glycerol, and 0.1 M phosphate buffer (PB)) and stored at −20°C before further analysis. After washing 4 times for 5 mins each in PB, 8-10 midbrain sections were then pre-incubated in 5% normal donkey serum and 0.3% Triton X-100 in 0.1 M PB for 30 minutes, and incubated overnight at 4 °C with the primary antibodies [rabbit anti-TH, 1:1000 (Thermo Scientific; Rockford, IL, USA); mouse anti-human α-synuclein 1:1000 (Thermo Scientific; Rockford, IL, USA); goat anti-luciferase 1:5000 (Abcam; Cambridge, UK)]. On the second day, sections were washed again and incubated for 1-hour with biotinylated species-specific secondary antibodies [donkey anti-rabbit/-mouse/-goat, 1:1000 (Jackson ImmunoResearch Laboratories Inc.; West Grove, PA, USA)], followed by 1-hour incubation in avidin-biotin-peroxidase solution (ABC Elite, Vector Laboratories; Burlingame, CA, USA). The protein was finally visualized by a 0.1M PB containing 5% 3.3’-diaminobenzidine (DAB) (Serva; Heidelberg, Germany) and 1% H_2_O_2_. The DAB-stained sections were mounted and counterstained with cresyl-violet and covered by mounting gel with coverslips (Corbit-Balsam; Kiel, Germany).

### 1.9. Double immunofluorescent labeling

The midbrain sections were washed for 20 minutes and pre-incubated in 10% normal donkey serum with 0.3% Triton X-100 in 0.1 M PB for 1 hour at room temperature (RT). Then they were incubated overnight at 4 °C with rabbit-anti-TH, 1:1000 (Thermo Scientific; Rockford, IL, USA) and mouse anti-human α-synuclein 1:1000 (Thermo Scientific; Rockford, IL, USA) or goat anti-luciferase 1:250 (Novus Biological; Littleton, CO, USA) primary antibodies followed by washing and incubation with fluorescence-labeled donkey anti-rabbit AlexaFluor488 and donkey anti-mouse or goat Cy3 secondary antibodies 1:500 (Jackson ImmunoResearch Laboratories Inc.; West Grove, PA, USA) for 2 hours at RT.

For microglial analysis, the same brain sections were incubated overnight with rabbit-anti-ionized calcium-binding adaptor (Iba1), 1:500 (Wako; Osaka, Japan) and sheep-anti-TH, 1:1000 (Merck Millipore; Darmstadt, Germany). On the next day, Iba1 positive (+) microglial cells were visualized with the biotinylated anti-rabbit secondary antibody followed by AlexaFluor647 conjugated Streptavidin and TH+ neurons were visualized with anti-goat AlexaFluor488. All epifluorescence images were acquired using an AxioImager M2 microscope (Carl Zeiss MicroImaging GmbH; Jena, Germany) equipped with an ORCA-Flash4.0 LT Digital CMOS camera (Hamamatsu C11440-42U; Hamamatsu City, Japan).

### 1.10. Unbiased Stereology and Optical density measures

The quantification of TH+ cells was performed by using Stereo Investigator software (v8, MicroBrightField, Magdeburg, Germany) at 40x magnification (Nikon Microphot-FX, Tokyo, Japan). The SNc was outlined as on every 5^th^ serial slice (2.4 to 4.1 mm dorsal of the bregma) and a fractionator probe was established for each section [28,37]. The criterion for counting an individual TH+ cell was the presence of its nucleus either within the counting frame or touching the upper and/or right frame lines (green), but not touching the lower and/or left frame lines (red). The number of TH+ nigral neurons was then determined by the Stereo Investigator program. The entire SNc was delineated from each animal in the groups of AN, AJ, LN, and LJ with a method described previously [39].

To quantify the intensity of microgliosis in the SNc and of TH immunoreactive fibers in the striatum, we measured the optical density (OD) of the slices. We compared the OD in the injected hemispheres to the non-injected hemispheres by FIJI software [40]. The slices were assessed in the identical anatomical coordinates described in the “quantification section” above. The striatum was outlined as previously described [41]. The striatum images were captured from three selected planes: +0.62 mm, +0.5 mm and +0.26 mm relative to bregma, and analyzed using Image J software version 1.43r for Mac OS X platform (http://rsbweb.nih.gov/ij/).

### 1.11. Statistical analysis

Statistical analysis was performed using the GraphPad Prism 7 (San Diego, CA, USA). Each data point represents mean ± S.E.M. \**p* < 0.05, \*\**p* < 0.01, \*\*\**p* < 0.001, \*\*\*\**p* < 0.0001 compared to control values. Multiple comparisons against a single reference group were performed by one-way ANOVA followed by a posthoc Tukey’s multiple comparisons test. For analysis of *in vivo* experiments, two-way ANOVA was performed to test the interaction of two variables: rAAV factor and JN403 treatment factor.

## 2. RESULTS

### 2.1. Establishment and characterization of αSyn toxicity using an in-vitro microglial model for evaluation of JN403

To determine whether JN403 elicits any cytotoxic effects in mouse primary microglial, we treated microglial cells with three different doses of JN403 (100 nM, 1 μM, and 10 μM) for 24 hours, followed by an assessment of cell viability using the MTT-assay. One-way ANOVA revealed the lack of cytotoxic effects of JN403 in concentrations up to 10 μM, and there was no difference between all these doses [F_3_, _4_ = 1.707, *p* > 0.05; **Fig. 2A**]. Based on this data, the two lower doses; 100 nM and 1 μM of JN403 were selected for all subsequent *in vitro* assays. Next, we verified the αSyn induced basal toxicity using 1μM of αSyn fragment AA 61-140 to provide a platform to examine the effects of JN403 on the level of proinflammatory factors in microglial cells. This αSyn fragment induced a significant reduction in the number of microglial cells. Therefore, 1μM of this αSyn fragment was applied for further experiments. Microglial cells were pre-treated with JN403 for 24 hours and then co-incubated with 1μM αSyn or control for another 24 hours. Cell viability was analyzed thereafter for a comparison of the JN403 treated to the control group without JN403. Treatment either with 100 nM or 1 μM of JN403 showed no rescue effect on cell viability. Data are presented as the mean ± standard error of the mean. Cell viability was measured for multiple independent experiments. Values for JN403 were found to be significantly different from the control group (\*\*\*\**p* < 0.0001) (**Fig. 2B**).

**Figure 2.**
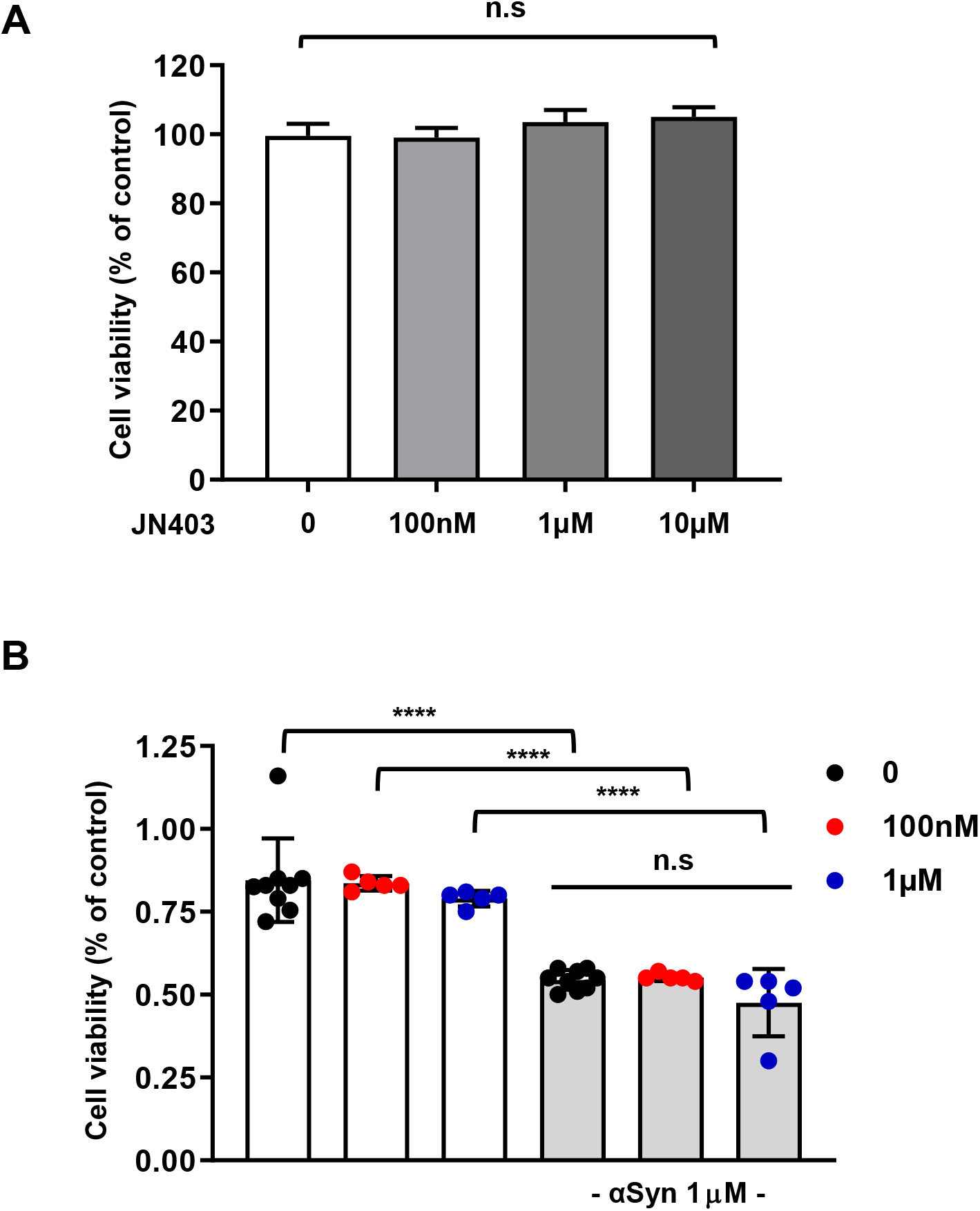
Establishment and characterization of αSyn toxicity using an in-vitro microglial model for evaluation of JN403. Microglial cells were incubated with three different doses of JN403 (100 nM, 1 μM, and 10 μM) for 24 hours, followed by an assessment of cell viability using the MTT-assay. (A) JN403 treatment alone did not elicit cellular toxicity in microglial cells at concentrations up to 1 μM. non-significant (n.s). (B) Microglial cells were pre-treated with 100 nM or 1 μM of JN403 for 24 hours and additionally coincubated for another 24 hours with 1μM αSyn fragment 61-140 on the next day. αSyn induced cytotoxicity in microglial cells. Neither 100 nM nor 1 μM of JN403 treatment changed the cell viability. Data are presented as the mean ± standard error of the mean. Cell viability was measured for multiple independent experiments. **** *p* < 0.0001, significantly different from the control group.

### 2.2 JN403 ameliorated αSyn induced NO and TNF-α release in microglial cell culture

Nitric Oxide (NO) has been considered as an inflammatory mediator and a neurotoxic effector molecule of immune responses [42,43]. We, therefore, selected NO as a parameter to elucidate the treatment effect of JN403. The microglial cells were pre-treated with JN403 for 24 hours but neither 100 nM nor 1 μM of JN403 induced NO release. Subsequently, 1μM αSyn was co-incubated with JN403 for another 24 hours to test the level of NO changes. When microglial cells were incubated with 1 μM of αSyn fragment alone, i.e. without JN403, NO release was dramatically increased. However, treatment either with of 100 nM or 1 μM of JN403 significantly decreased αSyn-induced NO release compared to the control [One-way ANOVA, (F_5_, _18_ = 76.68, (\*\*\*\**p* < 0.0001); **Fig. 3A**]. The results indicate that JN403 effectively downregulated the NO release in the αSyn model of microglial culture.

**Figure 3.**
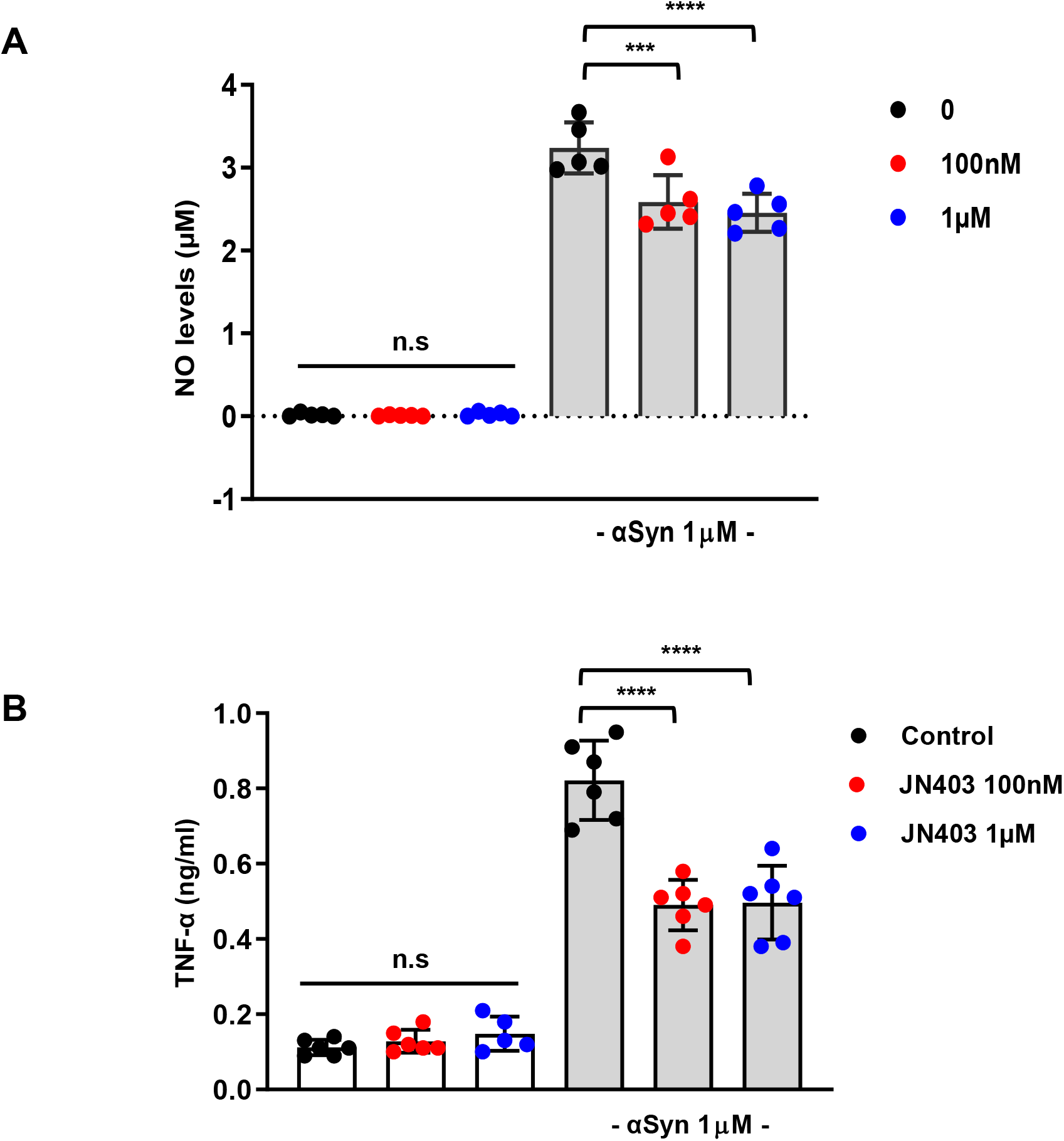
JN403 ameliorated αSyn induced NO and TNF-α release in microglial cell culture. Inflammatory parameters (A) Nitric Oxide (NO) and (B) TNF-α were measured in JN403 pre-treated microglial cells for a baseline. (A) NO, and (B) TNF-α release increased with 1μM αSyn incubation, but both 100 nM and 1 μM of JN403 co-treatment significantly reduced NO and TNF-α release, * *p* < 0.05, ** *p* < 0.01.

Next, changes in the cytokine release of TNF-α and IL-6 were quantified by ELISA in our microglial model. TNF is known to be essential for the complete expression of inflammation against external microbial invasion and the α7 subunit of nAChRs is known to be involved for cholinergic inhibition of TNF release [13,14,44]. On the other hand, IL-6 is known to play the role of both pro- and anti-inflammatory cytokine, which can modulate acute or systematic inflammation [45,46]. The ELISA results showed that TNF-α level was minimally increased during the pre-treatment of JN403 (100 nM or 1 μM) in microglial cells, but the difference between the control without JN403 exposure was statistically not significant. TNF-α level was significantly increased during the incubation with 1μM αSyn alone without JN403 treatment [One-way ANOVA, (F_5, 29_ = 98.97, \*\*\*\**p* < 0.0001); **Fig. 3B**]. Co-treatment, however, with either 100 nM or 1 μM of JN403 significantly decreased the TNF-α release compared to the control without JN403 (\*\*\*\**p* < 0.0001). Furthermore, the IL-6 level was also measured and found to be increased when 1μM αSyn was added to the microglia cells [F_5_, _18_ = 24.34, \*\*\*\**p* < 0.0001]. In contrast to the findings with TNF-α, neither 100 nM nor 1 μM of JN403 cotreatment altered IL-6 release in microglial cells compared to the control without JN403 (*p* > 0.05, data not shown). To conclude, the results demonstrated that JN403 lowered the pro-inflammatory mediators, especially the NO and TNF-α release, however not IL-6 in this αSyn fragment-activated microglial model, implying a specific role of JN403 to interact with the pro-inflammatory cytokine TNF-α.

### 2.3. Targeted human WT-αSyn overexpression via the delivery of recombinant AAV in the SNc

To test whether JN403 has the therapeutic potential to ameliorate the toxicity of overexpressed human αSyn on dopaminergic neurons or not, we employed the identical experimental design for the unilateral stereotactic operation targeting the SNc as previously published [28]. A recombinant adeno-associated virus (rAAV) harboring human wild-type αSyn (WT-αSyn) genome was unilaterally injected into the SNc to induce the αSyn overexpression. The control group received the rAAV vector harboring the genome for luciferase (luc). Three mice per group were selected 3 weeks post-injection to validate the transduction rate and double immunofluorescence staining against TH/αSyn or TH/luc was conducted with the midbrain slices. The percentage of the dopaminergic cells showing immunoreactivity against human αSyn or luc was calculated. Both viral vectors yielded high transduction rates in the dopaminergic SNc neurons of mice, with no significant difference between αSyn and luciferase transduction (91.7 ± 0.6% for αSyn and 93.4 ± 1.6% for luc). Immunoreactivity of αSyn was mainly observed in the injected hemisphere in comparison to the non-injected one 10 weeks post-injection (**Fig. 4A**). The immunohistochemistry staining shows that αSyn or luciferase was well expressed in most dopaminergic neurons in the SNc (**Fig. 4B-C**).

**Figure 4.**
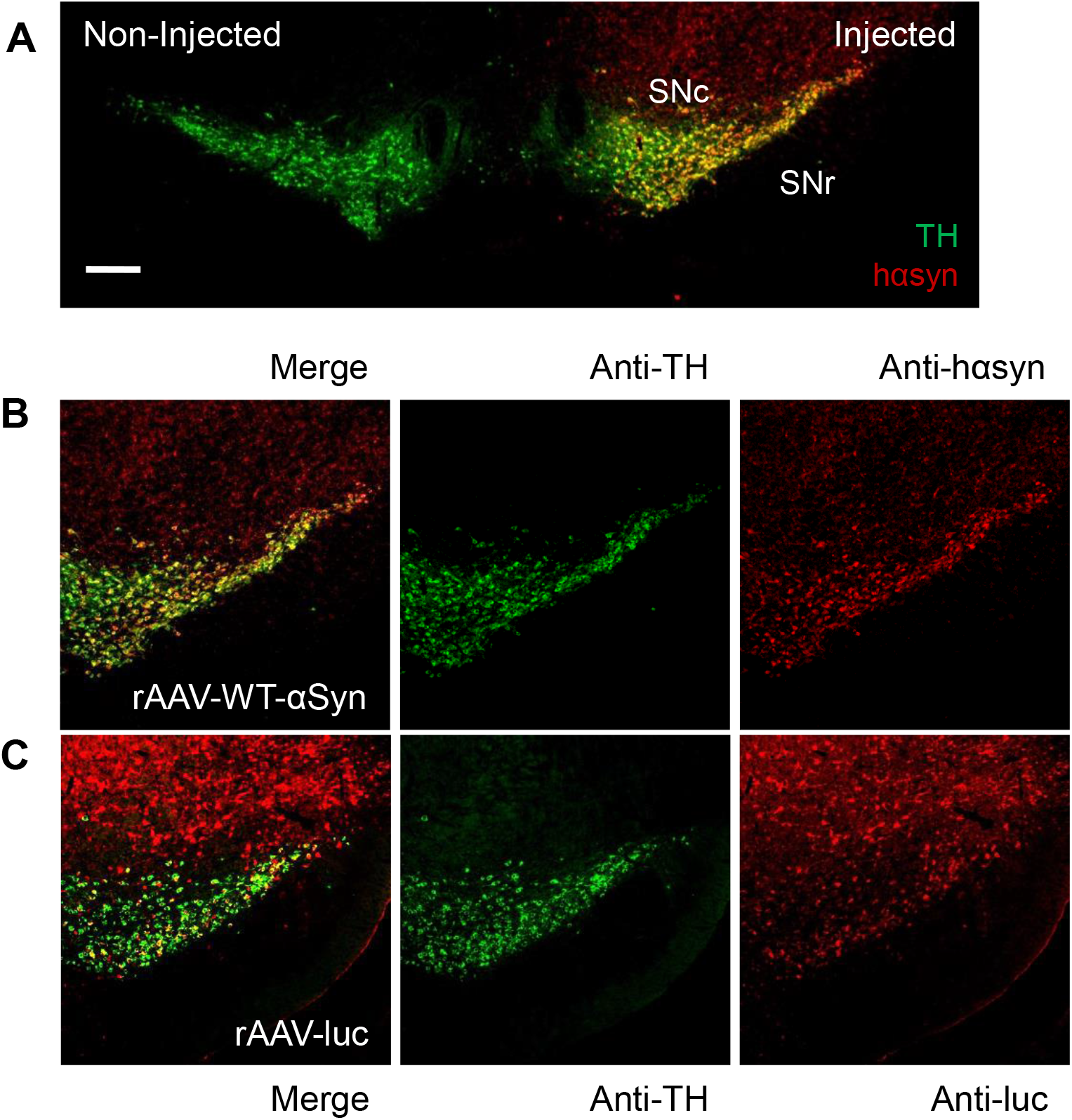
Targeted human WT-αSyn overexpression via the delivery of recombinant AAV in the SNc. Double immunofluorescence staining against TH/hαSyn or TH/luc was performed in midbrain slices 10 weeks after the stereotactic microinjection. (A) Merged image of TH (green) and hαSyn (red) showed that anti-hαSyn immunoreactivity is mostly seen in the injected side of substantia nigra compacta (SNc) with rAAV-WT-αSyn. Scale bar = 500 um. (B) Transduction of TH positive (+) nigral neurons with either rAAV-WT-αSyn or (C) rAAV-luc was confirmed. Anti-TH – green, anti-hαsyn or anti-luc – red. Scale bar = 200 μm.

### 2.4. No activated microglia observed from the nigral WT-αSyn overexpression model

To explore whether JN403 treatment regulates microglial activation *in vivo* after WT-αSyn or luc overexpression in the SNc, double immunofluorescent staining for TH and the microglial marker, Iba1 was performed. The representative photomicrographs of Iba1 fluorescent signals of all groups were explored and quantified accordingly. Histological results demonstrate that neither an increase in the number nor morphological change of microglia in the SNc was observed in both rAAV5-WT-αSyn and luc injected groups [F_1, 12_ = 2.678, (*p* > 0.05); n=4; **Fig. 5A-B**]. Moreover, the ratio of Iba1+ density from the injected compared to the non-injected side shows no significant differences between the rAAV groups and treatment [Two-way ANOVA; n=4, non-significant (n.s)]. For this reason, it was not feasible to test the efficacy of JN403 treatment on the microglial activation in this mouse model.

**Figure 5.**
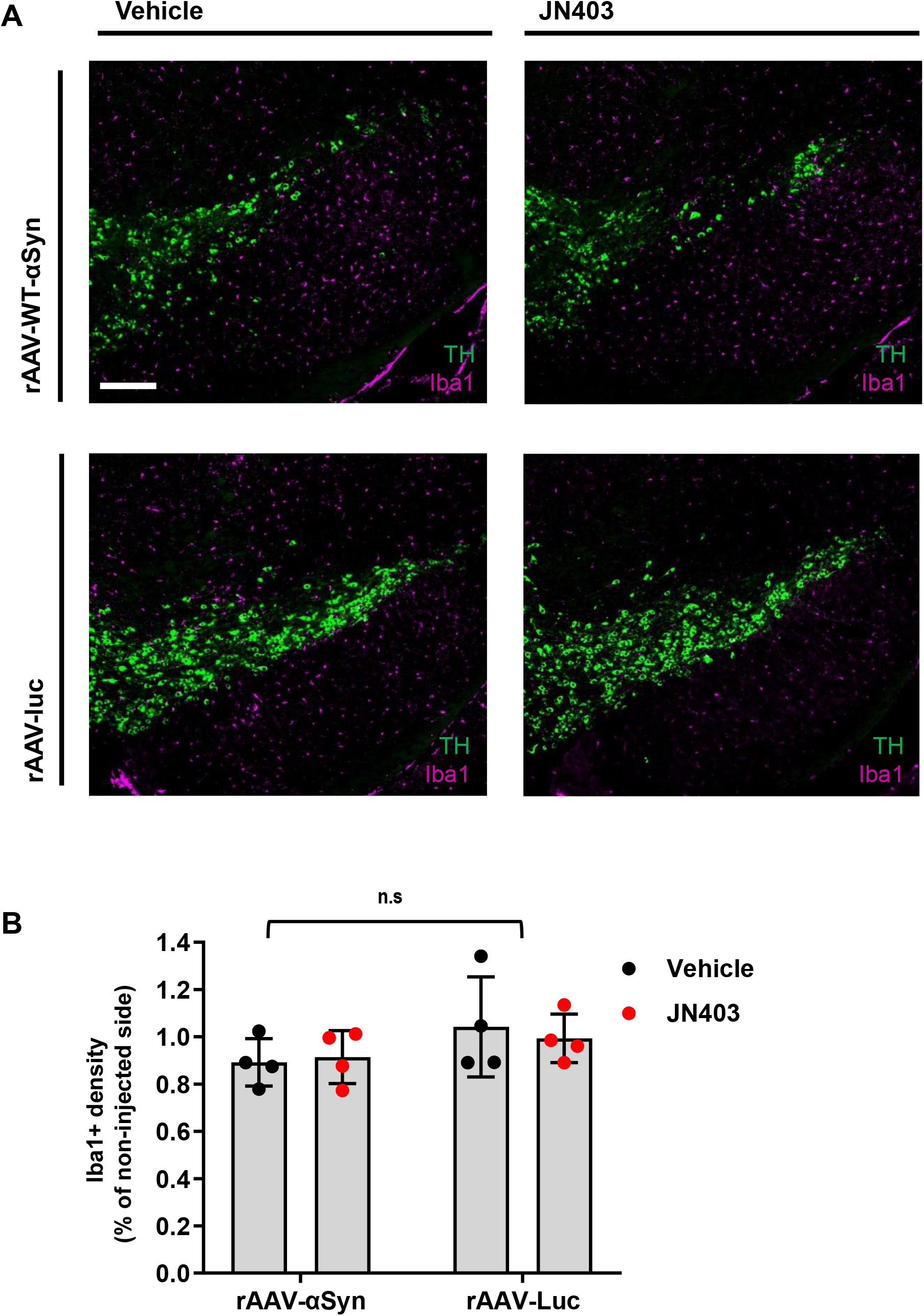
No activated microglia observed from the nigral WT-αSyn overexpression model. Iba1+ microglial cells were detected and the density analysis showed no differences between the rAAV groups and treatment groups. (A) Representative photomicrographs showing Iba1+ signals per group, TH (green) and Iba1 (magenta). No activated forms of microglia were found. Scale bar 200 μm. (B) The ratio of Iba1+ density of the injected side compared to the non-injected side showed no significant differences between the rAAVs and treatment, n=4, non-significant (n.s).

### 2.5. Lack of neuroprotective effect of JN403 on the reduction of TH+ neurons and nigrostriatal fibers

To investigate whether the number of TH+ neurons in the SNc remained unchanged, i.e. protected by JN403 in a mouse model of WT-αSyn overexpression, unbiased stereological quantification was performed. In the vehicle-treated rAAV-luc group, the number of TH+ neurons did not change [4526 ± 518.2 (non-injected) vs. 4440 ± 948.2 (injected side), *p* > 0.05; n=7], also in the group of JN403-treated rAAV-luc [4530 ± 741.3 (non-injected) vs. 4120 ± 877 (injected side), *p* > 0.05; n=6] indicating the microinjection or JN403 treatment per se did not reduce the number of TH+ neurons. In contrast, in the vehicle-treated rAAV-WT-αSyn group the number of TH+ neurons significantly decreased about 20% [4950 ± 600.3 (non-injected) vs. 3930 ± 566.2 (injected side), *p* < 0.05; n=8]. In addition, the JN403-treated rAAV-WT-αSyn group, also showed the reduced number of TH+ neurons [4563 ± 568.8 (non-injected) vs. 3710 ± 699.3 (injected side), *p* < 0.05; n=8, **Fig. 6A**]. The number of TH+ neurons in the injected side was expressed as a percentile compared to that of the non-injected side in all groups as shown in **Fig. 6B**. Two-way ANOVA analysis demonstrated that JN403 treatment did not influence the number of TH+ neurons in both rAAV-WT-αSyn and luc injected groups [F_1_, _25_ = 0.2556, (*p* > 0.05)]. A significant difference was only found between the two types of the rAAVs [F_1, 25_ = 9.068, (\*\**p* < 0.01)], as a result, there was no interaction between the rAAV and treatment factors (*p* > 0.05).

**Figure 6.**
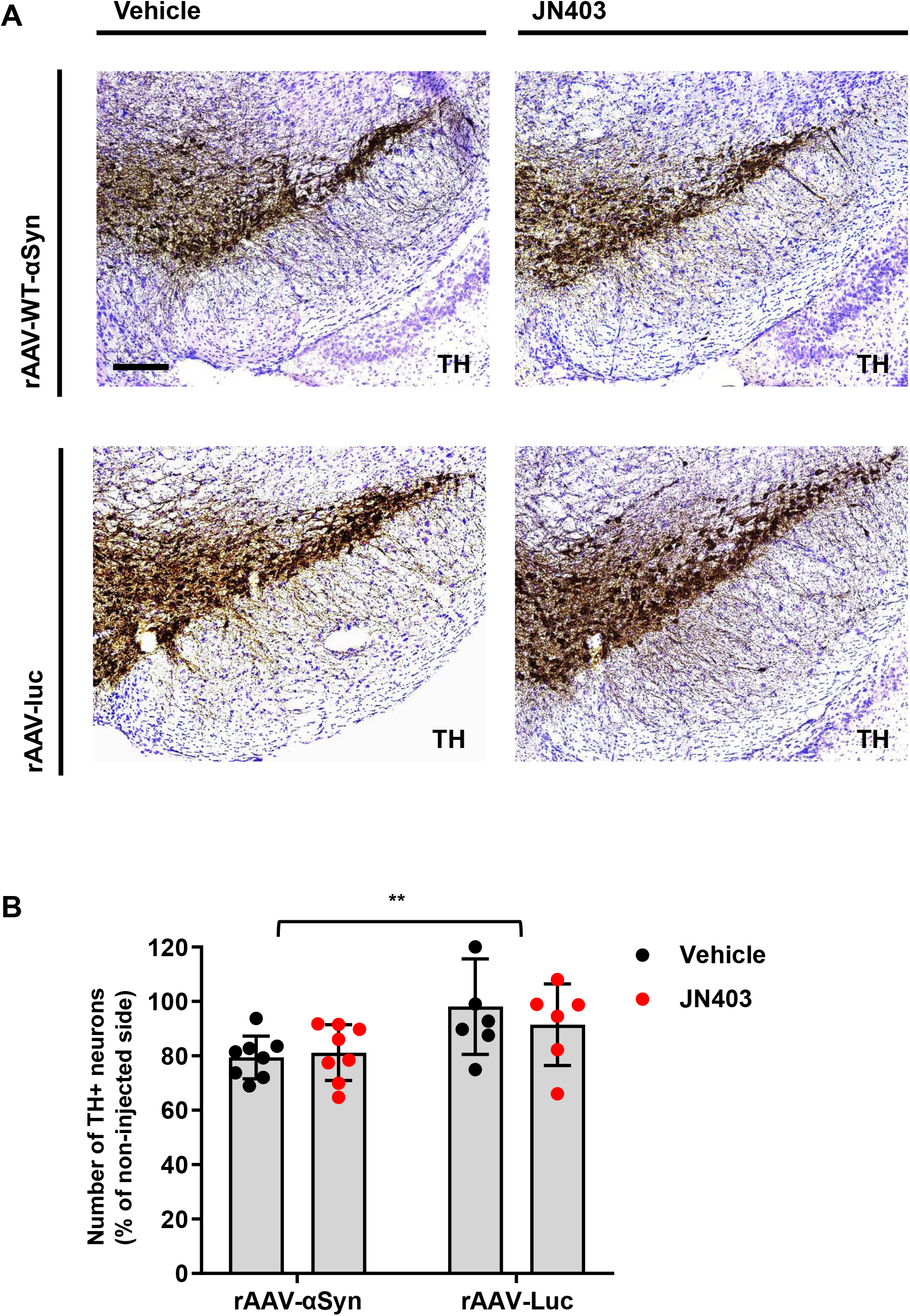
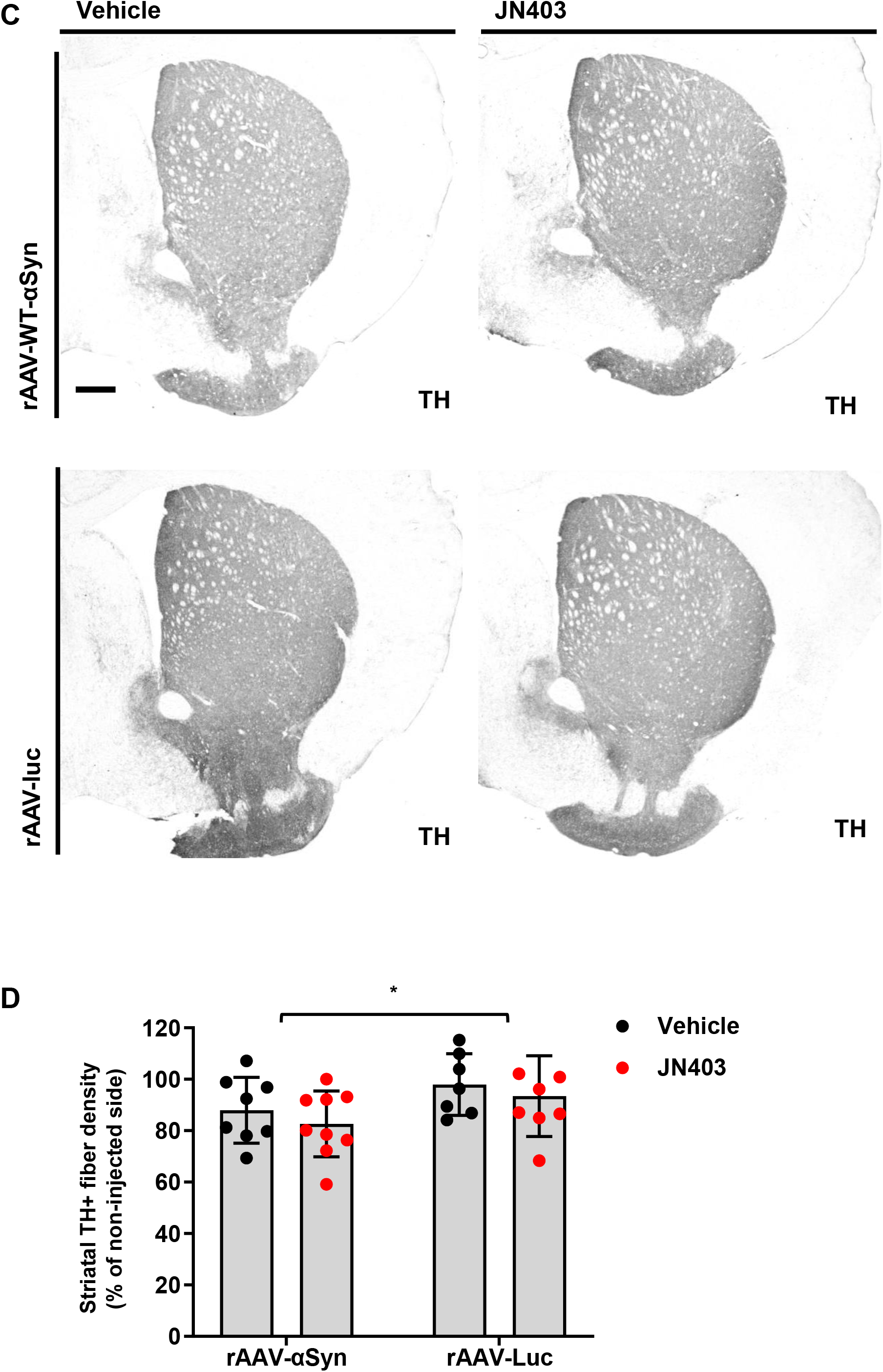
Lack of neuroprotective effect of JN403 on the reduction of TH+ neurons and nigrostriatal fibers. Stereological analysis of TH+ neurons in the SN showed no treatment effect. (A) Representative photomicrographs of TH+ nigral neurons in each group. Scale bar = 200 μm. (B) Plots show vehicle groups in black and JN403 treated groups in red for either rAAV-WT-αSyn or rAAV-luc injected animals. Percentage of TH+ nigral neurons in the injected compared to the non-injected side was analyzed; n=6-8, *p* **< 0.01 for rAAV factor; non-significant (n.s) for treatment factor. (C) Representative photomicrographs of the striatal TH+ staining 10 weeks after either intranigral rAAV-WT-αSyn or luc injection or JN403 or vehicle-treatment. Scale bar =500 um. (D) Optical density analysis of TH+ striatal fibers compared to the non-injected side. Plots in black show vehicle-treated and plots in red show JN403–treated groups in either rAAV-WT-αSyn or rAAV-luc injected animals; n=7-8, \**p* < 0.05 for rAAV factor.

To investigate the effect of WT-αSyn overexpression and JN403 treatment on the density of dopaminergic terminals, the optical density of TH+ fibers in the striatum was measured and the right side of regions (injected) were compared to the intact control side in each group. The two-way ANOVA analysis revealed that the striatal density of dopaminergic terminals was significantly reduced in the group of rAAV-WT-αSyn compared to the rAAV-luc injected groups [F_1, 28_ = 4.773, (\**p* < 0.05); n=7-8, **Fig. 6C-D**]. No JN403 treatment effect was observed and this result was consistent with the neuronal counting data [F_1, 28_ = 1.068, (*p* > 0.05)]. To conclude, treatment with JN403 did not alter the mean optical density of TH+ fibers in the group of rAAV-WT-αSyn, thus no significant difference was found in comparison to that of the vehicle-treated-control group.

## 3. DISCUSSION

In the current study, we investigated the anti-inflammatory effect of JN403, a novel α7-nAChR agonist in an *in vitro* microglial cell culture exposed to human αSyn fragment and a potential neuroprotective effect of JN403 on the survival of dopaminergic neurons in an *in vivo* mouse model using intranigral rAAV-mediated-WT-αSyn overexpression. Our results in microglial cell cultures showed that both 100 nM and 1 μM of JN403 decreased the release of NO and TNF-α, but not of IL-6 – as induced by the co-incubation with the synthetic αSyn fragment of AA 61-140. In the *in vivo* experiments, the primary target was to investigate the potential neuroprotective effect of JN403 on the survival of nigral dopaminergic neurons which were challenged by the focal rAAV-mediated overexpression of WT-αSyn or luc (as control) in mice. Unbiased stereology quantification demonstrated that overexpression of WT-αSyn in the SN reduced TH+ neuronal numbers by approximately 20 % compared to the non-injected side. Long-term daily subcutaneous application of JN403 - starting one week after stereotactic rAAV injection - did not protect the loss of dopaminergic neurons from αSyn-mediated toxicity in this mouse model. It was consistent with the results of the optical density analysis of the striatal dopaminergic terminals: JN403 was not able to revert the reduced density of the TH+ striatal fibers in the rAAV-WT-αSyn groups to the control level. In respect to a possible effect of JN403 on the activation of microglia in the SNc which may be associated with the overexpression of WT-αSyn, we failed to demonstrate a prominent microgliosis in the SNc. As a result, Iba1+ microglial density analysis showed no significant differences between all groups regardless of the JN403 administration.

To the extent of our knowledge, we propose the first microglial *in vitro* model evaluating the α7-nAChR agonist, JN403 using human αSyn fragment incubation, which is different from previously reported toxin-based experiments [18,47]. The αSyn fragment AA 61-140 at 1μM induced the most significant reduction in the number of microglial cells, compared to the other fragments. Therefore, this particular fragment was selected for the entire *in vitro* experiments to test inflammatory mediators in this study. Our *in vitro* findings in the inflammatory responses are in line with other recent studies, showing that pre-formed αSyn fibrils and purified total αSyn elevated the level of NO and TNF-α in primary mesencephalic culture or microglial cells [48–50]. However, one study reported that the IL-6 level was not significantly changed after the co-incubation with 200 nM αSyn in the microglial cell line that they employed [50]. The discrepancy may be due to the difference in the experimental design: we treated our primary microglial cultures with the commercially available αSyn fragment 61-140, whereas they incubated microglial cells with their own recombinant full-length αSyn. In our *in vitro* microglial cell culture model, JN403 exhibited a clear anti-inflammatory effect by reducing the αSyn-fragment induced the release of TNF-α and NO. These data are in agreement with other published results demonstrating that α7-nAChR agonists show concentration-dependent inhibition of the synthesis and release of NO and TNF-α [21,51,52]. The fact that JN403 did not show an effect on the IL-6 cytokine response in our microglial culture experiment may imply that IL-6 synthesis or release is not regulated by an α7-nAChR dependent pathway and therefore was not responsive to treatment with an α7-nAChR agonist such as JN403 [53]. However, we can not rule out other unknown molecular mechanisms of JN403 in the observed findings, that JN403 ameliorated increased level of the NO and TNF-α release, but not of the IL-6 release after αSyn fragment incubation.

For the *in vivo* study, we selected a PD model of intranigral rAAV-vector-mediated overexpression of WT-αSyn which induces a moderate loss of dopaminergic neurons. This model is – in principle - suitable for studies on a neuroprotective effect of a given compound, as in the employed PD model, the phosphodiesterase inhibitor vinpocetine was able to partially reverse this neuronal loss [28]. In the present setting, however, the *in vivo* part of our study had a major limitation: 10-weeks of WT-αSyn overexpression did not show microglial activation in the SNc. Thus this animal model did not provide an opportunity to investigate any potential matching mechanism (increase in proinflammatory cytokines) for the *in vivo* action of JN403, which was revealed in the *in vitro* part of this study. Moreover, respective biochemical measures of TNF, NO and IL-6 were not performed *ex vivo* from this animal model. One study on the Thy1-αSyn mice revealed that marked microglial activations as well as the increased TNF-α mRNA level and protein were first detected in the striatum at 1 month and in the SNc at 5-6 months of age [54]. This in-vivo finding demonstrates the differential correlation between the alpha-synuclein and TNF-α level per region at different time-windows both before and after nigrostriatal pathology. In our viral-mediated overexpression of αSyn model, we did not investigate an earlier time point to observe how microglial activation changes *in vivo* over time with the αSyn overexpression. However, unlike the Thy1-αSyn mice, we observed a 20 % loss of dopaminergic neurons in SNc as early as 10 weeks of viral overexpression in the absence of activated microglia. Our *in-vivo* result is therefore inconclusive but implies there may be distinctive molecular and pathological mechanisms beyond the dopaminergic degeneration, which remain to be further studied.

Another limitation of our *in-vivo* study design is that we did not measure the concentration of JN403 reached *in vivo* in the brain during the 9 weeks of experimental therapy. In the previous *in vivo* study JN403 (8.8 mg/kg) was orally administered in mice. The authors demonstrated, that that the concentration of JN403 in the brain was very similar to the compound’s EC50 (100 nM) at the α7-nAChR for at least 4 hours after the administration of JN403. Moreover, the high dose (30 mg/kg) of JN403 used in the current study was also evaluated and found to be tolerable in rats [32]. Our results do not exclude the possibilities of variances in the potency and pharmacokinetic profile in the class of α7-nAChR agonists. Therefore, at present, we can not distinguish whether the lack of a neuroprotective effect of JN403 on dopaminergic cell loss in the SNc is derived from our experimental design or rather the inefficacy of JN403.

Finally, it is important to point out, that all of the experimental evidence reporting on the positive effect of nicotine and α7-nAChR agonists has been obtained in mouse models of MPTP intoxication so far. In contrast, a few studies available - including our results herewith presented – generated in αSyn overexpression models have so far failed to demonstrate a beneficial effect of nicotine or an α7-nAChR agonist. This statement does not question the fact that the evidence for an acetylcholine-mediated reduction in neuroinflammation has accumulated. Thus enhancing this pathway may be beneficial to delay the pathogenesis of PD. It remains to be investigated whether α7-nAChR agonists would deliver such an anti-inflammatory effect at the nigral level and the other nuclei vulnerable to the etiopathogenesis of PD or even in the periphery such as the gastrointestinal system.

## 4. Conclusion

In this pilot study, we propose a novel *in vitro* model to study the effects of JN403, α-7-nicotine-acetylcholine-receptors (α7-nAChRs) agonist in human αSyn fragment exposed microglial cells. This suggests that JN403 reduces proinflammatory mediators, NO, and TNF-α, released upon αSyn incubation, and strategic targeting α7-nAChRs might be effective in ameliorating microglial activation. However, in a viral vector-mediated α-synuclein overexpression *in vivo* mouse model, no significant microgliosis was observed in both rAAV-WT-αSyn and luciferase injected groups, and JN403 treatment did not alter microglial density in the nigral regions of this model. Moreover, long-term in-vivo treatment of JN403 in this mouse model does not show a neuroprotective effect, thus failed to revert the reduction in the number of TH+ nigral neurons and TH+ striatal fiber density. Therefore, further research will be necessary using different PD-prodromal and PD-manifest animal models related to αSyn-etiopathogenesis to clarify whether and if so, in which part of the nervous system a long-lasting nAChRα7 activation may offer a PD-modifying effect.

## Acknowledgments

We truly thank Christine Höft for performing *in vitro* experiments and Sabine Anfimov for excellent histology expertise.

